# Genomic prediction by modelling genotype by environment interaction for yield in groundnut

**DOI:** 10.1101/2024.10.30.621054

**Authors:** Nelson Lubanga, Velma Okaron, Davis M. Gimode, Reyna Persa, James Mwololo, David K. Okello, Mildred Ochwo Ssemakula, Thomas L. Odong, Damaris A. Odeny, Diego Jarquin

## Abstract

Multi-environment trials (METs) are routinely conducted in plant breeding to capture the genotype-by-environment (G×E) interaction effects. We assessed the potential of using genomic prediction (GP) in four groundnut traits (pod yield[PY], shelling percentage[SP], seed weight[SW] and seed weight of 100 seeds[SW100]) observed in four environments. Three prediction models (M1: Environment + Line, M2: Environment + Line + Genomic, and M3: Environment + Line + Genomic + Genomic × Environment) were evaluated under four cross-validation (CV) schemes that simulate realistic problems that breeders face at different stages of their programs. CV2 predicts lines tested in sparse multi-location trials; CV1 predicts newly developed lines; CV0 predicts tested lines in unobserved environments by leaving one environment out; CV00 predicts untested lines in unobserved environments. Under CV2, the average predictive ability (PA) range was 0.6-0.68 (PY), -0.26-0.05 (SP), 0.5-0.59 (SW) and 0.79-0.86 (SW100). For CV1, the PA range was -0.12-0.51 (PY), -0.08-0.35 (SP), -0.10-0.47 (SW) and -0.05-0.47 (SW100). In CV0, the PA range was 0.27-0.35 (PY), -0.18--0.17 (SP), 0.24-0.32 (SW) and 0.59-0.66 (SW100). In CV00, the PA range was 0.01-0.23 (PY), -0.03-0.07 (SP), 0.01-0.19 (SW) and -0.02-0.38 (SW100). In all the four CV schemes, including marker data improved the PA. Incorporating G×E in M3 model improved PA in CV2 and CV1 schemes and also reduced the residual and environment variances. The high PA in CV2 imply that sparse testing could be implemented. The low PA recorded in CV00 shows that it is difficult to predict untested lines in new environments.

## 1. Introduction

Groundnut (*Arachis hypogaea L*.) is an important crop cultivated mainly by smallholder farmers in tropical and subtropical countries for food security and income generation. It is an important crop in sub-Saharan Africa (SSA), where it serves as a major source of dietary protein (Toomer, 2018), healthy fats (Mora-Escobedo et al., 2015), essential vitamins (Arya et al., 2016) and micronutrients (Kurapati et al., 2021). Despite its importance in SSA, groundnut yields remain extremely low with an average of ∼ 1017 kg/ha against a potential 4,505 kg/ha recorded in USA (FAO, 2022; USDA-NASS, 2021). This huge disparity makes breeding for high yielding groundnut varieties a key breeding objective (Asibuo et al., 2018; Dwivedi et al., 2003).

One of the significant threats to groundnut productivity in SSA is climate change that has resulted in heavy yield losses (Yeleliere et al., 2023), due to longer dry periods, heavy rainfall, hail damage, increased temperatures and emerging pests and diseases (Batley & Edwards, 2016). Developing climate-resilient groundnut varieties could significantly increase farmers yield and income through commercialization (Tabe-Ojong et al., 2023). One of the key steps in the varietal development process is multi-environment trials (METs) that help determine the genetic value of promising genotypes (Janila et al., 2013). METs help to assess the performance of genotypes across different environmental conditions, study genotype × environment (G×E) interaction and genotype stability, and predict the performance of untested genotypes (Burgueño et al., 2012). However, the presence of significant G×E interaction could potentially complicate the selection of high yielding groundnut genotypes in different environments (Asibuo et al., 2018).

Genomic selection (GS) is the application of genomic prediction (GP) models to estimate breeding values based on genome-wide markers in breeding programs (Meuwissen et al., 2001). The accurate capture of G×E helps to identify superior genotypes likely to perform better across different environments (Crossa et al., 2006). Single-environment GP models do not consider correlated environmental structures (Crossa et al., 2014), and therefore the presence of G×E is not accounted. This could lead to unreliable results (Crossa et al., 2017), due to the heterogeneity of genetic variance across environments, or the imperfect genetic correlation across environments (Lopez-Cruz et al., 2015). Mixed models that allow the incorporation of a genetic covariance matrix calculated from marker data, rather than assuming independence among genotypes could improve the estimation of the genetic effects (Burgueño et al., 2012; Jarquín, Crossa, et al., 2014). Modeling covariance matrices by incorporating G×E allows the use of information from correlated environments, which improves the accuracy of predicting genotypes (Burgueño et al., 2012). The benefit of using genetic covariance matrices in G×E mixed models is that the model relates genotypes across environments even when they are not present in all locations (Monteverde et al., 2018).

The first statistical framework for modeling G×E using genomic prediction was proposed by Burgueño et al. (2012), who extended the single-trait and single-environment genomic best linear unbiased prediction (GBLUP) model to a multi-environment context. They incorporated G×E interaction term in a GBLUP model using genetic and residual covariance functions. They found that the multi-environment GBLUP had a higher predictive ability than the single-environment GBLUP. However, they did not incorporate environmental variables to model G×E. Jarquín, Crossa, et al. (2014) introduced a reaction-norm model by modelling G×E as functions of molecular markers and environmental covariates. The main effects of molecular markers and environmental covariables, as well as their first order interactions (marker ×environmental covariates) are introduced using covariance structures that are functions of marker genotypes and environmental covariables. Lopez-Cruz et al. (2015) proposed a marker × environment interaction (M×E) GS model that can be implemented by regressing phenotypes on markers or using co-variance structures. Cuevas et al. (2016) extended the M×E model by using a non-linear (Gaussian) kernel to model G×E. Similarly, Crossa et al. (2016) also extended the M×E model using priors that produce shrinkage and variable selection i.e. Bayesian ridge regression (BRR) and BayesB (BB). In all these studies, the authors reported that models that incorporated G×E consistently had a higher predictive ability than single-environment GP models, regardless of the genetic correlation between environments.

In this study, we implemented three GP models to assess the predictive ability using four different cross-validation (CV) strategies that simulate unique realistic problems that breeders face at different stages of the breeding pipeline: CV2, predicting the performance of lines observed in some environments but not in others (sparse testing); CV1, predicting the performance of lines that have not been evaluated in any of the observed environments (newly developed lines); CV0, predicting tested lines in unobserved environments by leaving one environment out; CV00, predicting untested lines in unobserved environments (Jarquín et al., 2017). The main objectives of the present study were to 1) compare the performance of three prediction models in groundnut under four cross validation schemes that simulate different realistic problems that breeders face at different stages of the breeding pipeline, and 2) assess the impact of incorporating G×E on genomic predictive ability.

## 2. Materials and Methods

### 2.1. Phenotypic dataset

A total of 192 genotypes sourced from the International Crops Research Institute for the Semi-Arid Tropics (ICRISAT) were used in this study. ICRISAT utilises extensive germplasm collections to generate high yielding and climate-resilent varieties in Asia and Sub-Saharan Africa. The field trial evaluations were conducted in four locations: Serere (1.4994° N, 33.5490° E) and Nakabango (0° 19' 36” N, 33° 53' 36” E) in Uganda, and Chitala (13° 40' 59” S, 34° 16' 0” E) and Chitedze (13.9815° S, 33.6372° E) in Malawi. The trials were planted in an alpha-lattice design with two replications.

### 2.2. Phenotyping, genotyping and quality control

All the plant genotypes were harvested at the end of the growing season and evaluated for pod yield (PY) and other yield associated traits including shelling percentage (SP), seed weight (SW) and 100 seed weight (SW100) in the four environments. Details of the plant material, experimental design, phenotyping and genotyping procedure are described in Okaron et al. (2024). SNP filtering involved removing all markers with more than 20% missing values and a minor allele frequency below 5%. SNPs with a call rate lower than 80% and heterozygosity larger than 95% were also excluded. The final retained genotypes contained 38,853 SNPs. Filtered SNPs were converted to numeric allele classes (0, 1, 2) for homozygous minor, heterozygous and homozygous major alleles, respectively using Plink 1.9 (Purcell et al., 2007).

### 2.3. Phenotypic analysis

All the 192 genotypes observed in all the four environments (locations) were used in this study.The phenotypic analyses were performed in two stages. The first stage involved calculating the best linear unbiased predictors BLUPs of each genotype at each environment. The first stage was performed using ASReml-R Version 4.2 (Butler et al., 2023) in R statistical programming language (R Core Team, 2021) using the following model (equation 1):

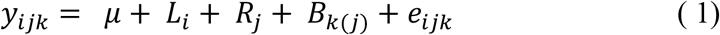

Here y_*ijk*_ represents the *i*^th^ genotype observed in the *j*^th^ replicate at the *k*^th^ incomplete block , *µ* is the overall mean (intercept), *L*_*i*_ is the random effect of the *i*^th^ genotype assuming 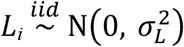, where N(.,.) denotes a normal density, *iid* stands for independent and identically distributed observations, and 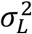 is the variance component of the genotypes; *R*_*j*_ is the fixed effect of the *j*^th^ replicate, *B*_*k*(*j*)_ is the random effect of the *k*^th^ incomplete block such that 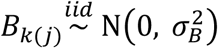, and 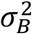 is the variance component of the block effect nested within replication; *e*_*ijk*_ is the random error term with 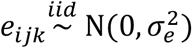 and 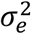 is the error variance addressing the non-explained variability by the previous model terms.

The broad sense heritability (*H*^2^) at each environment was calculated on an entry-mean basis and the genotypic and error variances were extracted from Equation 1. *H*^2^ was estimated as shown in equation 2:

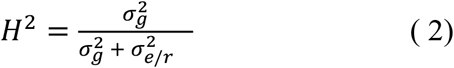

where 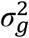 is genotypic variance, 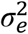 is error variance and *r* is the number of replicates within each environment.

### 2.4. Prediction models

The second stage of the analysis involved using the BLUPs obtained in the first stage (equation 1) to fit three different prediction models described below in equations 3, 5 and 6. The main aim of this study was to assess the performance of two genomic prediction models compared to a model based on phenotypic information only. The predictive ability of the three models under four different cross validation schemes were evaluated.

#### M1: Environment + Line (E + L)

In this model, the response variable corresponds to the adjusted phenotypes (*y*_*il*_) (BLUPs) obtained after fitting the corresponding mixed linear model as previously described. Consider that *y*_*ih*_ represents the response of the *i*^th^ genotype at the *h*^th^ environment. In this case, the environment and line effects are considered random outcomes:

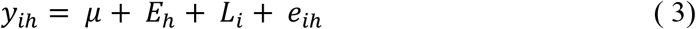

where *E*_*h*_ is the random effect of the *h*^th^ environment such that 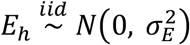 , and 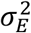 is the variance component of the environment effect, *L*_*i*_ is the random effect of the *i*^th^ line assuming 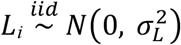, where 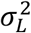 is the variance component of the genotype effect, and *e*_*ih*_ is the random error term, such that 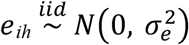 , where 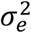 is the within-environment error variance.

#### M2: Environment, Line and Genomic main effects (E + L + G)

This model is an extension of Model 1 (equation 3). It considers the inclusion of the genomic random effect of the *i*^th^ genotype g_*i*_, which approximates its genetic value using molecular marker information. This model component can be defined by the regression on *p* marker covariates,

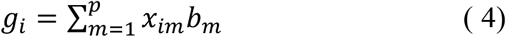

where *x*_*im*_ is the genomic information (marker matrix) of the *i*^th^ genotype at the *m*^th^ marker, and *b*_*m*_ is the corresponding marker effect, assuming 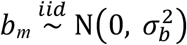 , and with 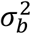 is the corresponding variance component which is homogenous to all markers. The vector of genomic effects **g =** {g_i_} follows a multivariate normal density with zero mean and variance-covariance matrix cov 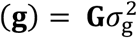 such that 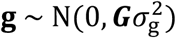 where 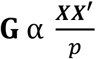 is the genomic relationship matrix, ***X*** is the centred and standardized (by columns) matrix of molecular markers and 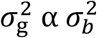 is the genomic variance. The resulting model becomes:

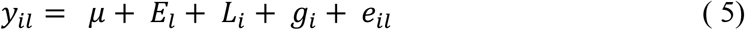

Where the genotype effects of the vector of genomic random effects **g** are correlated such that Model 2 allows the borrowing of information across genotypes. A disadvantage of this model is that the genomic estimated breeding values (GEBV) for a genotype tested in different environments is the same regardless of the environment. To allow a specific genomic effect in each environment, the interaction term (G×E) is included as shown in the Model 3.

#### M3: Environment, Line, Genomic and Genomic × Environment interaction effects (E+L+G+G×E)

This model is an extension of model 2 (Equation 5) by adding the interaction between the markers and environments (g*E*_*il*_) via covariance structures as described by Jarquín et al. (2014). This reaction norm model is an extension of the random effect Genomic Best Linear Unbiased Predictor (GBLUP) model where the interactions between each SNP and each environment are indirectly modelled using the following linear predictor:

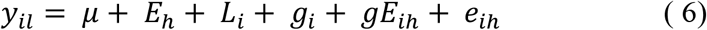

where the g*E*_*ih*_ term corresponds to the interaction between the genetic value of the *i*^th^ genotype and the *h*^th^ environment. This interaction term is assumed to follow a multivariate normal distribution such that 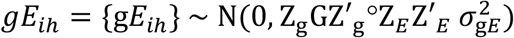 where, Z_g_ and Z_*E*_ are the correspondent incidence matrices for the effects of genetic values of genotypes and environments, respectively, 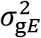 is the variance component associated to g*E* and '○' represents the Hadamard product (element-by-element product) between two matrices.

Jarquín et al. (2014) showed that assuming normality for the terms involving the interaction and also assuming that the interaction obtained using a first-order multiplicative model is distributed normally, then the covariance function is the Hadamard product of two covariance structures, one describing the genetic information and the other describing the environmental effects.

### 2.5. Model assessment under different cross-validation schemes

To assess the performance of the three different prediction models, four prediction strategies that are of interest for breeders were considered. These prediction scenarios attempt to mimic different realistic problems that breeders face at different stages of the breeding pipeline. A random 5-fold partitioning of the entire population was used for CV1 and CV2 and predictive ability was calculated as the average correlation between predicted and observed values of genotypes within the same environment for 10 random fivefold partitions. For CV0 and CV00, the correlations between observed and predicted values were computed for the leave-one-environment-out scenario as described by Persa et al. (2021). In all cases, correlations were computed between predicted and observed values within the same environment. Details of the four different cross validation scenarios are as described below:

a. CV2 (tested lines in observed environments) is mostly applicable when predicting incomplete field trials where some lines have been observed in some environments but not in others (e.g in sparse testing). The aim is to predict the performance of lines in environments where they have not yet been observed. In this study a random fivefold assignment was implemented where 20% of the phenotypic values were randomly assigned to each fold then one fold was used as testing set and the remaining four folds (80%) as training set for model calibration. This process was performed until all the five folds were completed. The procedure was repeated 10 times and the correlations between the predicted and observed values within the same environment were computed for all environments.
b. CV1 (untested lines in observed environments) is mainly used to predict the performance of newly developed lines that have not been observed in any of the tested environments, but there is available phenotypic information from other lines in these environments. The genetic similarities between genotypes in the training and testing sets influence the predictive ability of the model. In this strategy, lines are assigned to folds instead of phenotypes, such that all phenotypic records from the same lines are found in the same fold. Similar to CV2, a fivefold approach was implemented; however, the partitions were conducted on the lines (20% of the genotypes in each fold). Each fold was predicted using the phenotypic information of the remaining 4 folds (80% of the genotypes) used as training set. Each partition was predicted one at a time. The predictive ability was calculated as the correlations between predicted and observed values within the same environment and a total of 10 fivefold random partitions were performed.
c. CV0 (tested genotypes in unobserved environments) strategy relates the prediction of tested lines in unobserved environments, it is implemented by leaving one environment out. The performance of all the genotypes in one unobserved environment (one at a time) is predicted using the information of the remaining environments (training set). These steps are repeated for every environment. The predictive ability was evaluated as the correlations between the predicted and observed values for each environment.
d. CV00 (untested genotypes in unobserved environments) is similar to CV0 except that it also considers the prediction of novel genotypes that have not yet been tested in any environment. This strategy could be suitable in a situation where most of the genotypes are new or have not been observed before in any field and their performance for the planting season needs to be predicted. The accuracy of this strategy depends on the genetic and environmental similarities between genotypes in the testing set and those in the training sets.

All the prediction models were fitted using the BGLR package (Pérez & de los Campos, 2014).

## 3. Results

### 3.1. Phenotypic summary and broad sense heritability

Mean phenotypic values and broad sense heritabilities (*H*^*2*^) are provided for PY, SP, SW, and SW100 for the four environments in Table 1. The mean phenotypic values range was [46.97g-328.85g], [40.54%-64.06%], [22.73g-218.19g] and [28.08g-43.01g] for PY, SP, SW and SW100, respectively (Figure 1). Broad sense heritability ranged from [0.42-0.85], [0.00-0.79], [0.45-0.99] and [0.08-0.76] for PY, SP, SW and SW100, respectively (Table 1).

**Table 1.**
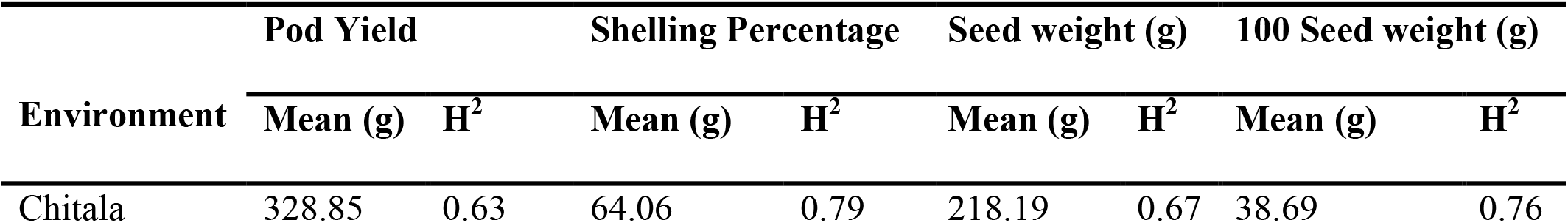
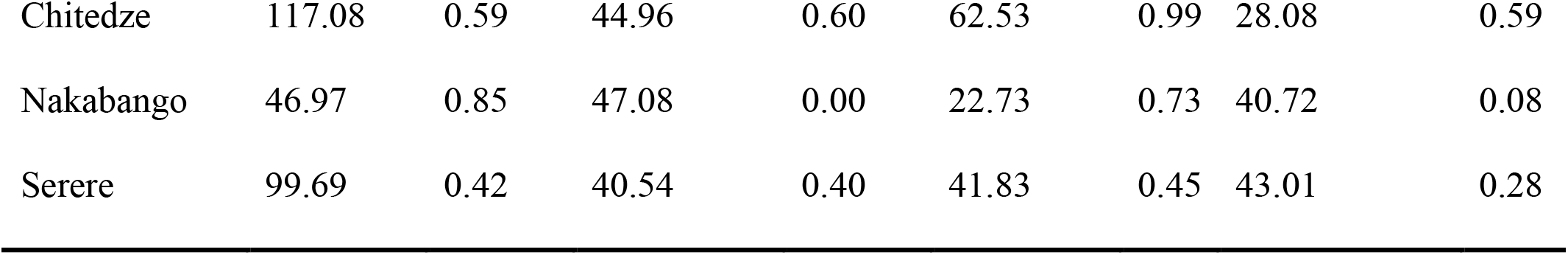
Distribution of phenotypic values and heritability of pod yield (PY), shelling percentage (SP), seed weight (SW) and 100 seed weight (SW100) in the four environments.

**Figure 1.**
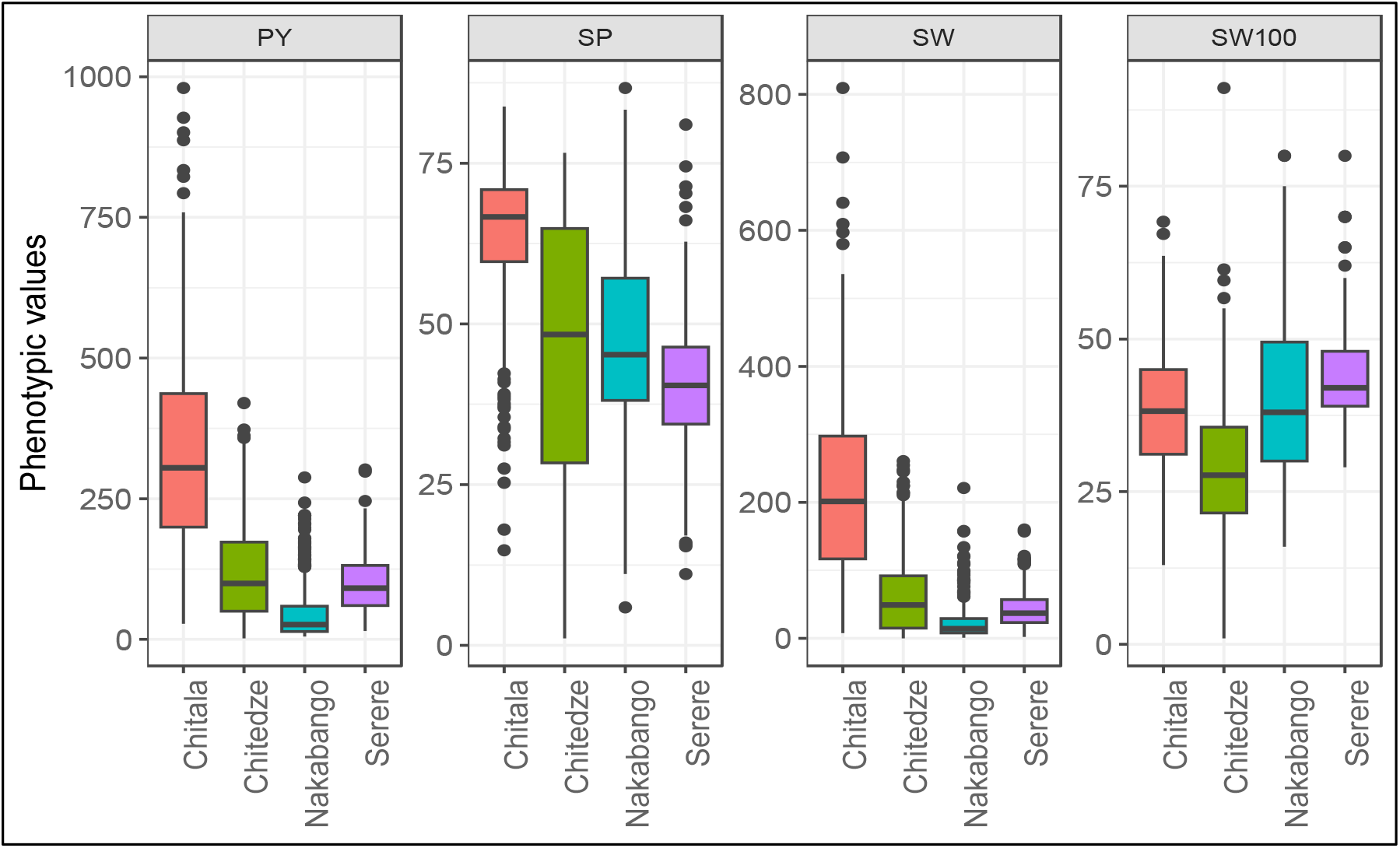
Box plot showing the distribution of pod yield (PY), shelling percentage (SP), seed weight (SW) and 100 seed weight (SW100) in the four environments.

**Figure 2.**
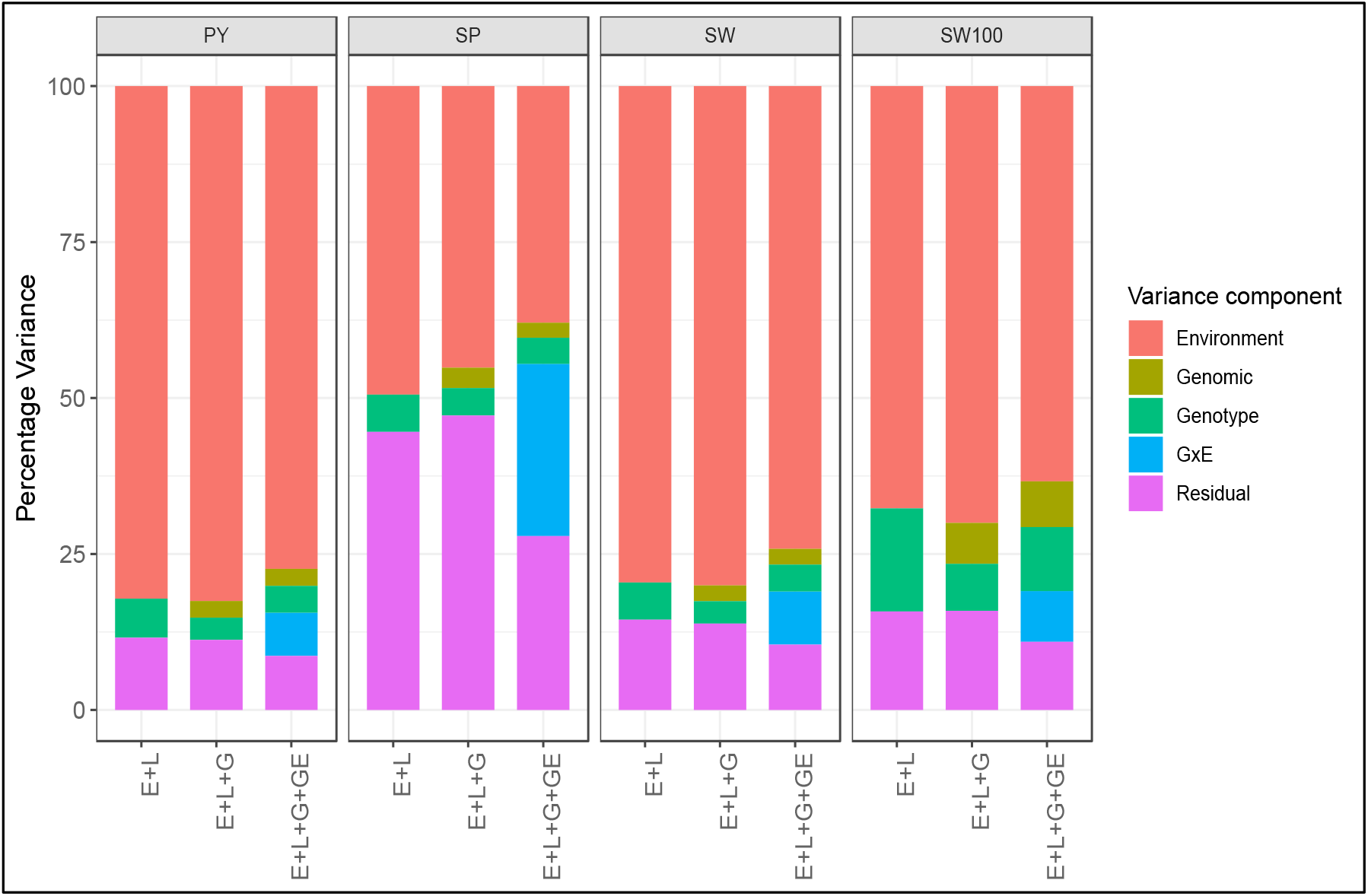
Percentage of explained variance components accounted for pod yield (PY), shelling percentage (SP), seed weight (SW) and 100 seed weight (SW100). The three models are M1 (E+L), M2 (E+L+G) and M3 (E+L+G+GE). E, the main effect of environments; L, the main effect of genotypes; G, the main effect of markers; and G×E, the interaction between genomic (markers) and environment.

### 3.2. Phenotypic correlations between the traits

Phenotypic correlations between traits showed that three traits were positively correlated i.e. PY, SP and SW. PY and SP had a correlation of 0.87, while PY and SW were highly correlated at 0.99. However as expected, there were no significant correlations between SW100 and the three traits that were positively correlated (PY, SP, SW). SW100 and PY had a correlation of 0.01, SW100 and SP showed correlation of 0.06, while SW100 and SW had a correlation of 0 (no correlation).

### 3.3. Variance Components

To better understand the sources of variability influencing the trait performance across environments, a full data analysis was conducted to compute the variance components.The relative percentage of variability explained by the different model terms for the three models M1 (E+L), M2 (E+L+G) and M3 (E+L+G+GE) is shown in Figure 3 for the four traits. The addition of G×E (interaction between genomic and environment) effects reduced the variability captured by the environment (E) and residual (R) terms. As expected, the main effects of environments consistently explained most of the total variance for all traits. The most complex model M3 (E+L+G+GE) had the lowest environment variance (E), while M1 had the highest in all the four traits. The estimated environment variance for PY was 82.2% (M1), 82.5% (M2) and 77.4% (M3). For SP, the environment variance was 49.5% (M1), 45.1% (M2) and 37.9% (M3). The environment variance for SW was 79.6% (M1), 80.0% (M2) and 74.4% (M3). For SW100, the environment variance was 67.7%, (M1), 70.0% (M2) and 63.4% (M3). Compared to M1, including the G×E term reduced the environment variance by 4.8%, 11.5%, 5.4% and 4.3% for PY, SP, SW and SW100, respectively. The G×E component explained 6.9%, 27.6%, 8.5% and 8.1% variation for PY, SP, SW and SW100, respectively (Figure 3).

Model M3 explained the lowest residual variance (R) in all the traits. Residual variance (R) calculated for PY was 8.7% (M3), 11.2% (M2) and 11.6% (M1). Residual variance (R) in SP was 27.9% (M3), 42.2% (M2) and 44.6% (M1). Residual variance in SW was 10.5% (M3), 13.9% (M2) and 14.5% (M1). For SW100, the residual variance was 10.9% (M1), 15.9% (M2) and 15.8% (M3). Similarly, when compared to M1, including the G×E term reduced the residual variance by 2.9%, 16.7%, 4.0% and 4.9% in PY, SP, SW and SW100, respectively (Figure 3).

### 3.4. Predictive ability in different cross-validation schemes

The mean predictive ability (PA) across 10 replicates and standard deviation (SD) for the three models i.e. M1 (E+L), M2 (E+L+G) and M3 (E+L+G+GE) estimated under each of the four cross-validation schemes (CV2, CV1, CV0 and CV00) for the four traits i.e. pod yield (PY), shelling percentage (SP), seed weight (SW) and 100 seed weight (SW100) are shown in Tables 2, 3, 4 and 5. In general, the predictive ability was higher in CV2 and CV00 compared to CV1 and CV0 for all the four traits. Here, the predictive ability was calculated by computing the average correlation between predicted and observed values within the same environment for all the 10 replicates. Incorporating the interaction between markers (genomic) and the environment (G×E) effects in model M3 increased predictive ability compared to the main effect models i.e. M1 (E+L) and M2 (E+L+G) for all traits except SW100 in CV2 and CV1.

**Table 2.**
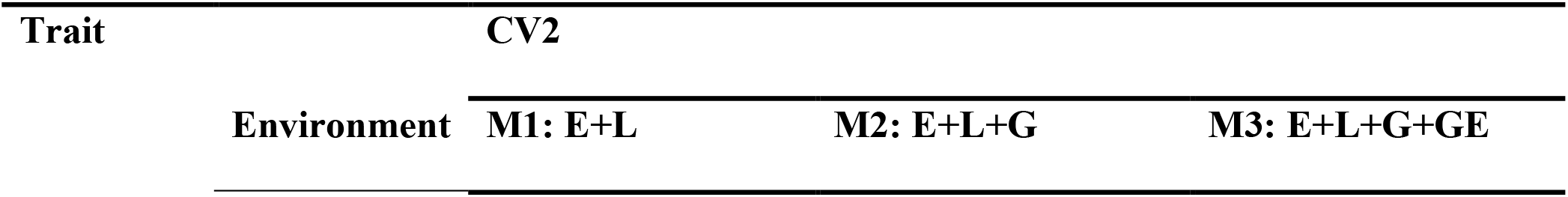
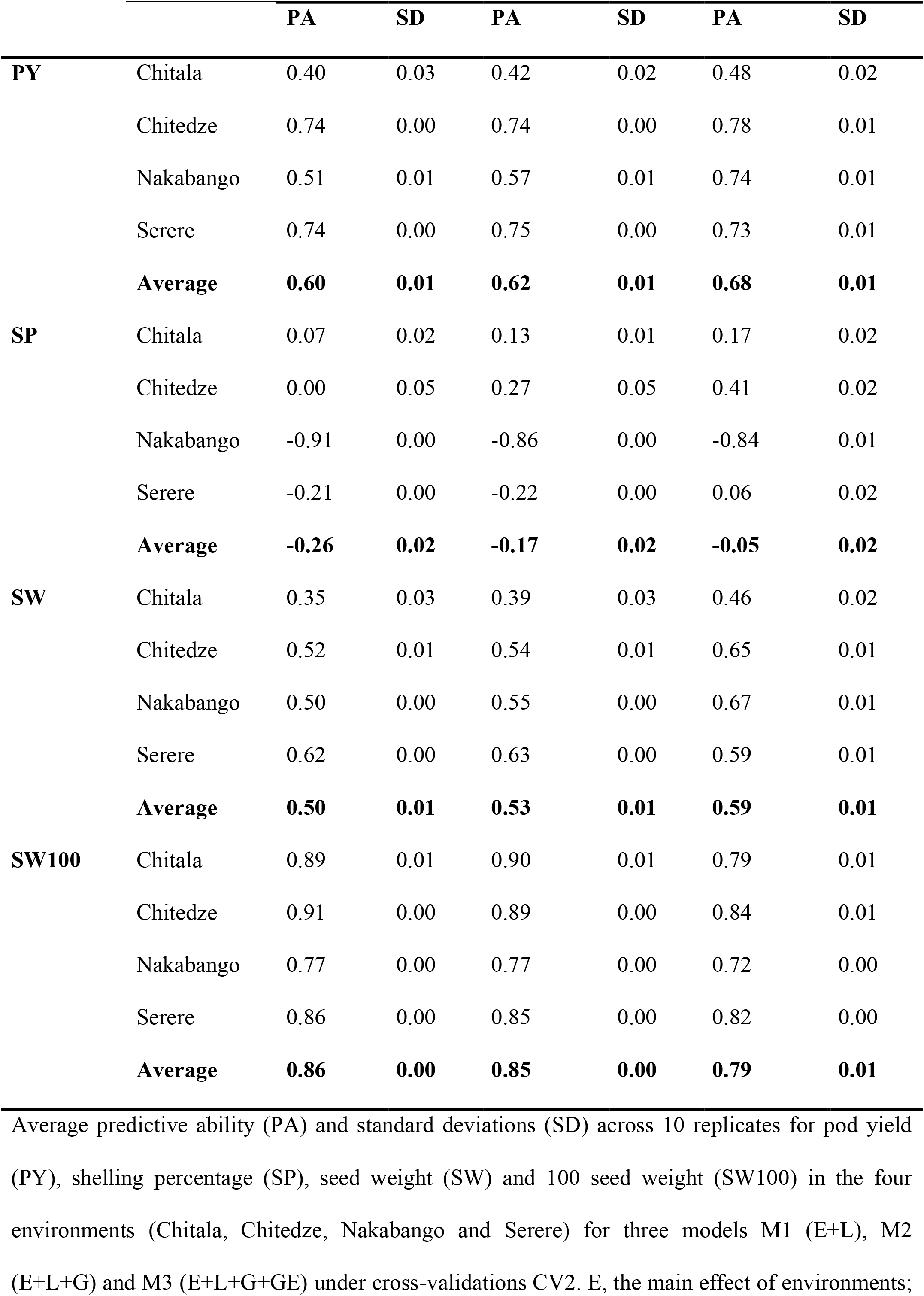
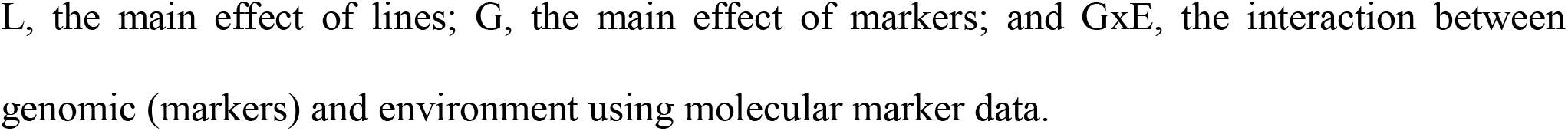
Predictive abilities under CV2 (tested genotypes in tested environments i.e. sparse testing)

**Table 3.** Predictive abilities under CV1 scenario (untested genotypes in tested environments i.e. newly developed lines)

| Trait | Environment | CV1 |  |  |  |  |  |
| --- | --- | --- | --- | --- | --- | --- | --- |
|  |  | M1: E+L |  | M2: E+L+G |  | M3: E+G+L+GE |  |
|  |  | PA | SD | PA | SD | PA | SD |
| PY | Chitala | -0.15 | 0.05 | 0.28 | 0.03 | 0.37 | 0.02 |
|  | Chitedze | -0.13 | 0.08 | 0.43 | 0.02 | 0.49 | 0.03 |
|  | Nakabango | -0.11 | 0.07 | 0.54 | 0.01 | 0.67 | 0.01 |
|  | Serere | -0.08 | 0.07 | 0.48 | 0.02 | 0.49 | 0.02 |
|  | **Average** | **-0.12** | **0.07** | **0.44** | **0.02** | **0.51** | **0.02** |
|  | Chitala | -0.14 | 0.05 | 0.16 | 0.04 | 0.19 | 0.02 |
|  | Chitedze | -0.12 | 0.03 | 0.32 | 0.04 | 0.42 | 0.02 |
| SP | Nakabango | 0.02 | 0.07 | -0.34 | 0.02 | 0.42 | 0.06 |
|  | Serere | -0.09 | 0.07 | -0.19 | 0.02 | 0.36 | 0.03 |
|  | **Average** | **-0.08** | **0.05** | **-0.01** | **0.03** | **0.35** | **0.03** |
|  | Chitala | -0.14 | 0.04 | 0.28 | 0.03 | 0.38 | 0.02 |
|  | Chitedze | -0.15 | 0.05 | 0.32 | 0.03 | 0.45 | 0.03 |
| SW | Nakabango | -0.09 | 0.07 | 0.47 | 0.02 | 0.64 | 0.01 |
|  | Serere | -0.06 | 0.06 | 0.38 | 0.02 | 0.43 | 0.02 |
|  | **Average** | **-0.11** | **0.06** | **0.36** | **0.02** | **0.47** | **0.02** |
|  | Chitala | -0.11 | 0.08 | 0.53 | 0.02 | 0.55 | 0.01 |
|  | Chitedze | -0.09 | 0.09 | 0.47 | 0.01 | 0.48 | 0.01 |
| SW100 | Nakabango | 0.00 | 0.07 | 0.42 | 0.01 | 0.40 | 0.03 |
|  | Serere | -0.01 | 0.08 | 0.46 | 0.01 | 0.45 | 0.02 |
|  | **Average** | **-0.05** | **0.08** | **0.47** | **0.01** | **0.47** | **0.02** |

Average predictive ability (PA) and standard deviations (SD) across 10 replicates for pod yield (PY), shelling percentage (SP), seed weight (SW) and 100 seed weight (SW100) in the four environments (Chitala, Chitedze, Nakabango and Serere) for three models M1 (E+L), M2 (E+L+G) and M3 (E+L+G+GE) under cross-validations CV1. E; the main effect of environments; L, the main effect of lines; G, the main effect of markers; and GxE, the interaction between genomic (markers) and environment using molecular marker data

**Table 4.** Predictive ability under CV0 scheme (tested genotypes in untested environments)

|  |  | CV0 |  |  |  |  |  |
| --- | --- | --- | --- | --- | --- | --- | --- |
|  |  | M1: E+L |  | M2: E+L+G |  | M3: E+L+G+GE |  |
| Trait | Environment | PA | SD | PA | SD | PA | SD |
| PY | Chitala | 0.31 | 0.09 | 0.38 | 0.06 | 0.37 | 0.06 |
|  | Chitedze | 0.29 | 0.17 | 0.43 | 0.16 | 0.23 | 0.12 |
|  | Nakabango | 0.24 | 0.15 | 0.27 | 0.13 | 0.22 | 0.10 |
|  | Serere | 0.26 | 0.14 | 0.32 | 0.12 | 0.26 | 0.11 |
|  | **Average** | **0.27** | **0.14** | **0.35** | **0.12** | **0.27** | **0.10** |
| SP | Chitala | 0.07 | 0.03 | 0.15 | 0.03 | 0.10 | 0.05 |
|  | Chitedze | 0.02 | 0.07 | 0.04 | 0.06 | -0.03 | 0.03 |
|  | Nakabango | -0.61 | 0.13 | -0.67 | 0.14 | -0.64 | 0.16 |
|  | Serere | -0.14 | 0.05 | -0.20 | 0.06 | -0.15 | 0.06 |
|  | **Average** | **-0.17** | **0.07** | **-0.17** | **0.07** | **-0.18** | **0.08** |
|  | Chitala | 0.33 | 0.06 | 0.37 | 0.03 | 0.37 | 0.04 |
|  | Chitedze | 0.20 | 0.13 | 0.30 | 0.13 | 0.13 | 0.09 |
| SW | Nakabango | 0.24 | 0.15 | 0.29 | 0.15 | 0.23 | 0.10 |
|  | Serere | 0.24 | 0.12 | 0.30 | 0.11 | 0.23 | 0.12 |
|  | **Average** | **0.25** | **0.12** | **0.32** | **0.10** | **0.24** | **0.09** |
|  | Chitala | 0.32 | 0.09 | 0.51 | 0.20 | 0.46 | 0.12 |
|  | Chitedze | 0.82 | 0.05 | 0.84 | 0.02 | 0.82 | 0.07 |
| SW100 | Nakabango | 0.57 | 0.13 | 0.59 | 0.11 | 0.64 | 0.05 |
|  | Serere | 0.65 | 0.10 | 0.70 | 0.09 | 0.66 | 0.10 |
|  | **Average** | **0.59** | **0.09** | **0.66** | **0.10** | **0.64** | **0.09** |

Predictive ability (PA) and standard deviations (SD) for pod yield (PY), shelling percentage (SP), seed weight (SW) and 100 seed weight (SW100) in the four environments (Chitala, Chitedze, Nakabango and Serere) for three models M1 (E+L), M2 (E+L+G) and M3 (E+L+G+GE) under cross-validations CV0. E, the main effect of environments; L, the main effect of lines; G, the main effect of markers; and GxE, the interaction between genomic (markers) and environment using molecular marker data

**Table 5.** Predictive ability under CV00 scheme (untested genotypes in untested environments)

|  |  | CV00 |  |  |  |  |  |
| --- | --- | --- | --- | --- | --- | --- | --- |
|  |  | M1: E+L |  | M2: E+L+G |  | M3: E+L+G+GE |  |
| Trait | Environment | PA | SD | PA | SD | PA | SD |
| PY | Chitala | -0.05 | 0.10 | 0.17 | 0.04 | 0.15 | 0.04 |
|  | Chitedze | 0.00 | 0.09 | 0.22 | 0.12 | 0.18 | 0.12 |
|  | Nakabango | 0.05 | 0.08 | 0.26 | 0.11 | 0.21 | 0.14 |
|  | Serere | 0.03 | 0.07 | 0.28 | 0.14 | 0.15 | 0.13 |
|  | **Average** | **0.01** | **0.08** | **0.23** | **0.10** | **0.17** | **0.11** |
| SP | Chitala | -0.04 | 0.07 | 0.12 | 0.05 | 0.04 | 0.10 |
|  | Chitedze | -0.02 | 0.08 | 0.04 | 0.08 | 0.07 | 0.06 |
|  | Nakabango | -0.04 | 0.07 | -0.26 | 0.09 | -0.12 | 0.09 |
|  | Serere | 0.02 | 0.07 | -0.20 | 0.06 | -0.13 | 0.11 |
|  | **Average** | **-0.02** | **0.07** | **-0.07** | **0.07** | **-0.03** | **0.09** |
| SW | Chitala | -0.04 | 0.08 | 0.14 | 0.03 | 0.12 | 0.04 |
|  | Chitedze | -0.01 | 0.08 | 0.14 | 0.10 | 0.12 | 0.09 |
|  | Nakabango | 0.04 | 0.08 | 0.25 | 0.10 | 0.18 | 0.14 |
|  | Serere | 0.03 | 0.06 | 0.22 | 0.11 | 0.10 | 0.10 |
|  | **Average** | **0.01** | **0.08** | **0.19** | **0.09** | **0.13** | **0.09** |
|  | Chitala | -0.05 | 0.06 | 0.34 | 0.14 | 0.29 | 0.13 |
|  | Chitedze | -0.03 | 0.06 | 0.46 | 0.02 | 0.43 | 0.03 |
| SW100 | Nakabango | -0.01 | 0.05 | 0.34 | 0.07 | 0.34 | 0.08 |
|  | Serere | 0.02 | 0.03 | 0.39 | 0.07 | 0.35 | 0.08 |
|  | **Average** | **-0.02** | **0.05** | **0.38** | **0.08** | **0.35** | **0.08** |

Predictive ability (PA) and standard deviations (SD) for pod yield (PY), shelling percentage (SP), seed weight (SW) and 100 seed weight (SW100) in the four environments (Chitala, Chitedze, Nakabango and Serere) for three models M1 (E+L), M2 (E+L+G) and M3 (E+L+G+GE) under cross-validations CV00. E; the main effect of environments; L, the main effect of lines; G, the main effect of markers; and GxE, the interaction between genomic (markers) and environment using molecular marker data.

#### 3.4.1. Predictive ability under CV2 (predicting the performance of genotypes observed in some environments but not in others (sparse testing approach)

Table 2 shows the results of the average predictive abilities under the incomplete field trial scenario (CV2). For PY model M3 (0.68) had the highest average predictive ability followed by M2 (0.62) and M1 (0.60). The predictive ability for PY ranged from 0.40-0.74 (M1), 0.42-0.75 (M2) and 0.48-0.0.78 (M3). SP was the most difficult trait to predict under this scenario as it had the lowest predictive ability for all the models and traits. The average predictive ability was -0.05, -0.17 and -0.21 for M1, M2 and M3 models, respectively. The predictive ability for SP ranged from -0.91-0.007 (M1), -0.86-0.27 (M2) and -0.84-0.41 (M3). For SW, the average predictive ability was 0.59, 0.53, 0.50 for model M3, M2 and M1, respectively. The predictive ability for SW ranged from 0.35-0.62 (M1), 0.39-0.63 (M2) and 0.46-0.67 (M3). SW100 recorded the highest predictive ability in all the three models compared to PY, SP and SW. The average predictive ability was 0.86, 0.85 and 0.79 for M1, M2 and M3, respectively. The predictive ability for SW100 ranged from 0.77-0.89 (M1), 0.77-0.90 (M2) and 0.72-0.82 (M3) (Table 2).

#### 3.4.2. Predictive ability under CV1 (predicting the performance of lines that have not been evaluated in any of the observed environments)

Table 3 presents the average predictive ability for the CV1 cross-validation scenario. Here, model M3 had the highest predictive ability, while M1 had the lowest for all the traits. The average predictive ability were -0.12 (M1), 0.44 (M2) and 0.51 (M3) for PY. The predictive ability range was -0.15--0.08 (M1), 0.28-0.54(M2) and 0.37-0.69(M3). The average predictive ability for SP were -0.08 (M1), -0.01(M2) and 0.35 (M3). The predictive ability ranged from -0.14-0.02 (M1), -0.34-0.32 (M2) and 0.19-0.42 (M3). For SW, the average predictive ability was -0.11 (M1), 0.36 (M2) and 0.47 (M3). The predictive ability for SW ranged from -0.15--0.06 (M1), 0.28-0.47(M2) and 0.38-0.64 (M3). The average predictive ability for SW100 was -0.05 (M1), 0.47 (M2) and 0.47 (M3). The predictive ability for SW100 ranged from -0.11-0.00 (M1), 0.42-0.53 (M2) and 0.40-0.55(M3) (Table 3) (Table 3).

#### 3.4.3. Predictive ability under CV0 (predicting tested lines in unobserved environments by leaving one environment out)

Table 4 presents the results of the scenario of mimicking the prediction of observed genotypes in novel environments. Here, the inclusion of G×E in M3 did not lead to improvement in predictive ability compared to M2 and M1 for all the traits (Table 3). For instance, the average predictive ability for PY was 0.27 (M1), 0.35 (M2) and 0.27 (M3). The predictive ability range for PY was 0.24-0.31 (M1), 0.27-0.43 (M2) and 0.22-0.37 (M3). SP was the most difficult trait to predict under CV0 and the average predictive ability was -0.17 (M1), -0.17(M2) and -0.18 (M3). The predictive ability range for SP was -0.61-0.07 (M1), - 0.67-0.15(M2) and -0.64-0.10 (M3). The average predictive ability for SW was 0.25 (M1), 0.32 (M2) and 0.24 (M3). The predictive ability range for SW was 0.20-0.33 (M1), 0.29-0.37 (M2) and 0.13-0.37 (M3). For SW100, the average predictive ability was 0.59 (M1), 0.66 (M2) and 0.64 (M3). The predictive ability range was 0.32-0.82 (M1), 0.51-0.84 (M2) and 0.46-0.82 (M3) (Table 4).

#### 3.4.4. Predictive ability under CV00 (predicting new genotypes in novel environments)

Table 5 presents the predictive ability for the most challenging breeding scenario (CV00), where model M2 performed slightly better than M3 for all the traits except SP (Table 5). The average predictive ability for PY was 0.01 (M1), 0.23 (M2) and 0.17 (M3). The predictive ability range for PY was -0.05-0.05 (M1), 0.17-0.28 (M2) and 0.15-0.21 (M3). Similarly, SP was the most difficult trait to predict under this scenario and the average predictive ability was -0.02 (M1), -0.07 (M2) and -0.02 (M3). The predictive ability for SP ranged from -0.04-0.02 (M1), -0.26-0.12 (M2) and -013-0.07 (M3). The average predictive ability for SW was 0.01 (M1), 0.19 (M2) and 0.13 (M3). The predictive ability range for SW was -0.04-0.03 (M1), 0.14-0.25 (M2) and 0.10-18 (M3). For SW100, the average predictive ability was -0.02 (M1), 0.38 (M2) and 0.35(M3). The predictive ability range for SW100 was -0.05-0.02 (M1), 0.34-0.46 (M2) and 0.29-0.43 (M3) (Table 5).

## 4. Discussion

In this section, we discuss the results obtained by implementing three genomic prediction models M1 (E+L), M2 (E+L+G) and M3 (E+L+G+GE) in groundnut METs under four cross validation schemes. We discuss the variance components and predictive ability of the three models for pod yield (PY), shelling percentage (SP), seed weight (SW) and 100 seed weight (SW100).

### 4.1. Variance components

The main challenge in the analysis of multi-environment trials (MET) in plant breeding and variety testing is modelling and exploiting genotype–by-environment interaction (Crossa et al., 2016). It is important to estimate reliable and accurate prediction of genotype's performance across environments in crop improvement programs. The use of single-environment models could be unreliable in the presence of significant genotype-by-environment interaction (Burgueño et al., 2012; Cuevas et al., 2016), due to the heterogeneity of genetic variance across environments, or imperfect genetic correlation of the same traits across sites/seasons (Cuevas et al., 2016). In this study, we implemented the reaction norm model M3: E+L+G+GE (Jarquín, Crossa, et al., 2014), which accounts for the interaction between molecular markers and the environment (G×E) through covariance structures. Then a comparison of performance of the model M3 versus two main effects models i.e. model M2(E+L+G) and M1(E+L) was conducted. The relative contribution of each model term was studied through the computation of variance components.

The addition of the G×E interaction in M3 substantially decreased the amount of variance captured by both the environment and residual terms in all the four traits as compared to the traditional genomic prediction model (M2) and the phenotypic selection model M1. Compared to the traditional GP model (M2), including the G×E term reduced the environment variance by 5.1% (PY), 7.2%(SP), 5.9%(SW) and 6.7%(SW100) and residual variance by 2.6% (PY), 19.3% (SP), 3.4% (SW) and 5.0%(SW100). Model M3 (E+L+G+GE) explained the lowest environment (E) and residual variance (R) in all the four traits. This imply that the inclusion of G×E term increases the proportion of variance accounted for by the model and could improve the predictive ability. Canella Vieira et al. (2022) reported that the model incorporating G×E decreased the amount of variability captured by the environment and residual terms as compared to the model that did not include this interaction term in soybean. Similarly, Mageto et al. (2020) reported that when marker information was incorporated in the prediction model (*i.e*., M2 and M3), the estimated variance due to environments was substantially reduced in maize populations. These authors also reported that including the interaction term (genomic × environment) in the prediction model reduced the residual variance component by ∼30% in model M3.

### 4.2. Predictive ability under different cross validation schemes

Genomic prediction is a modern approach that uses genome-wide markers to predict the genetic merit of un-phenotyped individuals and can reduce the breeding cycle time and increase the selection accuracy. Compared to the traditional phenotypic selection, GS can improve the rate of genetic gain and accelerate the release of new improved varieties in breeding programs by: 1) reducing the generation interval since new parents can be selected at the earliest seedlings stage rather than in late field testing stages, 2) increasing the accuracy of selecting superior genotypes at the seedlings stage where selection accuracy is generally low, 3) increasing the selection intensity at the seedlings stage by testing more seedlings at reduced cost through sparse testing, 4) shortening the entire breeding cycle time by reducing or eliminating the lengthy field testing steps, while recycling parents with high breeding value at early generations back into the breeding program. Genomic prediction is currently implemented in breeding programmes in low to middle income countries (LMIC) with limited resources e.g. cassava (Okeke et al., 2017; Wolfe et al., 2017), tea (Lubanga et al., 2023) and pearl millet (Srivastava et al., 2020). Cross-validation (CV) schemes are useful in genomic prediction in assessing the prediction accuracy for different traits and environments (Burgueño et al., 2012; Jarquín et al., 2017). Four different cross-validation (CV) schemes simulating different realistic problems that breeders face were used in this study i.e. CV2, CV1, CV0, CV00 (Jarquín et al., 2017; Persa et al., 2021). As expected, a higher predictive ability was obtained in CV2 and CV00 compared to CV1 and CV0 scenarios for all the four traits. This is because all the lines have already been observed in at least one environment and therefore there is already more information available that is included in the genomic relatedness and correlated environments in the calibration set. However, in CV1 and CV00, prediction is performed for genotypes that have not been evaluated previously (Jarquín et al., 2017). In general, SP were the most difficult trait to predict. This can be seen from the low predictive ability observed in all the CV schemes for all the three models. This could be because SP had the highest GxE, residual and environmental variance compared to all the other traits. SP had the lowest heritability (0.00) at Nakabango, which could also imply possile errors in the precision of data collection process at this site.

Under CV2 scheme (predicting the performance of genotypes observed in some environments but not in others i.e. in sparse testing), the model M3 (E+L+G+GE) had the highest predictive ability compared to M2 (E+L+G) and M1 (E+L) for all the traits except SW100. This implies that incorporating the genomic × environment (G×E) improved predictive ability in sparse testing. The model M3 and M2 had a significantly higher predictive ability compared to M1, indicating that using molecular marker information is beneficial in sparse testing. Similar findings were also observed by Mageto et al. (2020) in maize and Canella Vieira et al. (2022) in soybean, who reported higher prediction accuracies in M3 compared to models that did not include G×E. In the CV2, information from related lines (including the targeted line tested in other environments) and correlated environments (including the targeted environment) is used, and prediction assessment benefits from borrowing information between lines within an environment, between lines across environments, and among correlated environments (Burgueño et al., 2012). This is reflected in the high predictive ability results observed for all the traits under CV2 compared to other CV schemes.

In CV1 (predicting the performance of lines that have not been evaluated in any of the observed environments (i.e. newly developed lines), M3 (E+L+G+G×E) similarly showed the highest predictive ability compared to M2 (E+L+G) and M1 (E+L) for all the traits except SW100. In both CV2 and CV1 schemes, the model M3 had higher predictive ability in all the traits except SW100. This CV scheme is more challenging than predicting the performance of lines that have already been evaluated in different but correlated environments (e.g. under CV2 scheme). A similar observation was reported by Burgueño et al. (2012), Jarquín et al. (2017) and Canella Vieira et al. (2022), who reported higher predictive abilities in CV2 scheme compared to CV1. This highlights the value of incorporating information from correlated environments when predicting the performance of lines. Selection of lines without field testing, as simulated in CV1, allows shortening of the generation interval, but the predictive ability is much lower than in CV2, which might compromise the annual rate of genetic gain in a GS breeding program.

In CV0 (predicting tested lines in unobserved environments by leaving one environment out), there was no improvement in predictive ability by including the G×E component since the performance of model M3 (E+L+G+G×E) was similar or less than that of M2 (E+L+G) and M1 (E+L). This could be because most lines have already been observed in some environments and therefore could be considered as replicates in the unobserved environments. It could also imply that there is plenty of information already available for most lines, hence good predictions in the missing environments. Similar findings were observed by Jarquín et al. (2017) who observed that including G×E did not improve predictive ability in CV0 in a Kansas wheat breeding program.

CV00 (predicting new genotypes in novel environments) is the most difficult breeding scenario to predict. All the three models under CV00 had the lowest predictive ability, implying that the performance of models is improved when there is available information from closely related genotypes and environments that could be borrowed (incorporated into the model). However, there was no benefit of including the G×E component since the performance of model M3 (E+L+G+G×E) was similar or less than that of M2 (E+L+G) and M1 (E+L).

### 4.3. Sparse testing of groundnut in multi-environment trials

In our study, sparse testing scenario i.e. predicting the performance of genotypes observed in some environments but not in others was simulated under CV2. All the three models i.e. M3 (E+L+G+G×E), M2 (E+L+G) and M1 (E+L) had a higher predictive ability under CV2 for all the traits compared to other CV schemes i.e. CV1, CV0 and CV00. Similarly, the model incorporating genomic × environment interaction (G×E) had the highest predictive ability in all the traits. Similar findings were also reported by Mageto et al. (2020) in maize, Canella Vieira et al. (2022) in soybean and Jarquín et al. (2017) in wheat. They reported that all the models under CV2 had higher predictive ability compared to CV1, CV0 and CV00. Modeling genetic and residual covariances improved the predictive ability in CV2 compared to other schemes. Modeling G×E allows borrowing of information within lines and across environments. This is not possible in CV1 and CV00 for instance because the lines being predicted have not been evaluated in any environment. This implies that sparse testing using a model incorporating G×E could be a good strategy for evaluating groundnut genotypes in METs.

Sparse testing is a prediction strategy that optimizes the allocation of resources in which the phenotypic information of lines is split across environments (Jarquin et al., 2020). This implies that not all genotypes are tested in all environments due to the high cost of setting up METs and the limited resources such as land, water, time and seed availability. The best way to benefit from sparse testing is to establish METs that are cost effective, without negatively impacting the selection accuracy of genotypes in breeding trials. According to Jarquin et al. (2020), the accuracy of predicting unobserved genotypes in METs is influenced by (a) the number of overlapping (OL) genotypes across environments, (b) the number of environments where each genotype is tested, (c) the implemented prediction model, and (d) the number of phenotypes per genotype (Garcia-Abadillo et al., 2024). Further work to evaluate the efficiency of different sparse testing designs should be conducted to assess their effectiveness in groundnut breeding programs.

## 5. Conclusion

Genotype × environment interaction (G×E) is one of the key issues affecting breeding value estimation accuracy for agronomic traits in plant breeding. Including the G×E interaction term in the GP model reduced the environment and residual variances. Additionally, including G×E in M3 improved predictive ability under CV2 and CV1 scenarios. Higher predictive ability was observed in CV2 and CV0, compared to CV0 and CV00, implying that borrowing information from environments by modelling the variance– covariance matrix is beneficial for predicting new genotypes. Sparse testing could be implemented to improve the selection accuracy and reduce the cost of conducting METs in groundnut breeding programs.

## 6. Data availability

The R scripts and the phenotypic and genomic datasets used to perform the analysis are publicly available on the figshare repository: https://figshare.com/articles/dataset/Datasets_and_R_scripts_used_to_perform_Genomic_prediction_in_multi-environment_groundnut_traisl/27324150?file=50058195

## 7. Acknowledgements

The first author is thankful to the staff of the International Crops Research Institute for the Semi Arid Tropics (ICRISAT) in Malawi and the national staff in Uganda who helped in the data collection. This work was largely supported by the Food and Agriculture Organisation (FAO) through the benefit sharing fund of the International Treaty on Plant Genetics for Food and Agriculture. Supplemental funding was provided by the Scholarship Advert for Short Term Academic Mobility (SCIFSA) and the Regional Universities' Forum for Capacity Building in Agriculture (RUFORUM) to the second author.

